# Warming and species richness weaken eco-phenotypic feedback loop in long-term natural ecosystems

**DOI:** 10.1101/2025.03.23.644810

**Authors:** Qinghua Zhao, Yingying X. G. Wang, Chi Xu, George Sugihara, Frederik De Laender

## Abstract

Eco-phenotypic feedback—reciprocal interactions between phenotypic traits and ecological dynamics—is increasingly recognized as a driver of biodiversity patterns, species interactions, and ecosystem functioning. Through this feedback, phenotypic traits such as body size can rapidly respond to environmental variation through plastic or evolutionary changes, altering population abundance, which in turn feeds back to shape the trait dynamics. Yet, whether the integrity of this feedback remains stable under environmental change remains unclear. Using long-term monitoring data from 101 aquatic and terrestrial ecosystems encompassing multiple generations, we provide the first synthesis showing that warming and species richness systematically weakens the eco-phenotypic feedback loop, while mean species body size strengths it. Our findings reveal that climate change can erode key trait– demography couplings and highlight the importance of integrating ecophenotypic frameworks into global change research.

## Introduction

Environmental change can rapidly alter both phenotypic traits and population abundance (Gardner *et al*. 2011; McLean *et al*. 2016). For instance, warming can reduce body size through increased metabolic costs while lowering population density via reduced survival or recruitment (Gardner *et al*. 2011); similarly, predation often favor smaller-bodied individuals while directly decreasing abundance through higher mortality (Bergquist *et al*. 1985; Wellborn 1994). More broadly, changes in phenotypic traits and population abundance are not independent but often interdependent (Gibert *et al*. 2022; Govaert & Klauschies 2024; Han *et al*. 2023). Trait shifts can modify demographic rates, influencing population density, which in turn feeds back to shape trait dynamics through plasticity or selection (Govaert & Klauschies 2024). When these reciprocal influences occur together, they form a coupled loop in which ecological and phenotypic changes reinforce and modify each other over time (Govaert & Klauschies 2024; Han *et al*. 2023). This coupled dynamic gives rise to eco-phenotypic feedback (Govaert & Klauschies 2024; Han *et al*. 2023).

Unlike eco-evolutionary feedback that requires heritable genetic changes, eco-phenotypic feedback can operate through both heritable genetic changes and nonheritable mechanisms, including behavioral shifts, and physiological adjustments (Govaert & Klauschies 2024). Eco-phenotypic feedback has been documented in a range of taxa and environments, from single species to food webs, regulating population persistence, trophic interactions, community structure, and ecosystem resilience under ongoing environmental changes (Gibert *et al*. 2022; Govaert & Klauschies 2024; Han *et al*. 2023). For instance, in fish populations subject to intense harvesting, reduced body size can lower fecundity and recruitment, driving population decline; lower densities then relax intraspecific competition for resources, potentially allowing body size to increase in subsequent generations (Conover & Munch 2002; Hutchings & Reynolds 2004; Law 2000). Similarly, in mussel beds, high density increases competition for planktonic food and space, limiting growth and reducing average shell size (Bayne & Worrall 1980; Okamura 1986). Smaller mussels have reduced fecundity, lowering recruitment and abundance over time (Bayne & Worrall 1980; Gosling 2003). Lower density then reduces competition and increases resource availability, enabling larger bodied shell growth in surviving individuals and recruits (Bayne & Worrall 1980; Okamura 1986). This feedback shapes mussel bed structure, competitive interactions with other sessile invertebrates, and the resilience of intertidal communities (Rietkerk *et al*. 2021; Van De Koppel *et al*. 2008).

However, strength of eco-phenotypic feedback is not fixed but context dependent, shifting with environmental conditions (Govaert & Klauschies 2024; Han *et al*. 2023). Climate warming—a pervasive anthropogenic driver—can alter both trait dynamics and population abundance (McLean *et al*. 2016; Schwarz *et al*. 2017). Rising temperatures can influence phenotypic traits through metabolic constraints, developmental acceleration, and shifts in life history strategies (McLean *et al*. 2016). Concurrently, warming can modify population abundance by affecting reproductive rates, mortality, dispersal, and interspecific interactions (Parmesan 2006; Walther *et al*. 2002). Although numerous studies have examined these effects independently, few have explicitly investigated how warming disrupts their coupling, with existing evidence largely coming from experiments and no studies from natural ecosystems. Beyond temperature, eco-phenotypic feedback strength may also be shaped by biodiversity such as species richness. Greater richness increases the likelihood of diverse interspecific interactions—competition, facilitation, predation—that can alter both traits and population abundance trajectories (Tilman 1999). Trait variation within populations, such as mean body size, can further buffer or amplify the feedback strength; species with smaller mean size often exhibit shorter generation times and greater plasticity, potentially tightening or weakening the feedback (Kingsolver & Pfennig 2007; Peters 1983).

However, no study has explicitly documented the effect of species richness and mean body size on the integrity of eco-phenotypic feedback. We address these gaps by conducting a synthesis evaluating the ecophenotypic feedback integrity under warming, biodiversity and trait variation in natural ecosystems. We compiled long-term, continent-wide data from 101 ecological systems spanning aquatic and terrestrial environments, including plankton, fish, and ground beetle communities, spanning 10 to 47 years and encompassing multiple generations per species and enabling robust detection of trait–phenotypic feedback. Specifically. We use Convergent Cross Mapping (CCM), a model-free technique for inferring causal relationships, to quantify bidirectional feedback between the phenotypic trait (body size) and population dynamics (Fig 1). We then assess how the prevalence of this feedback varies with climate warming and biotic drivers such as species richness and average trait values. By linking eco-phenotypic feedback to key abiotic and biotic drivers, this study strengthens the conceptual and empirical basis for incorporating eco-phenotypic feedback into ecological theory and forecasting.

**Figure 1.**
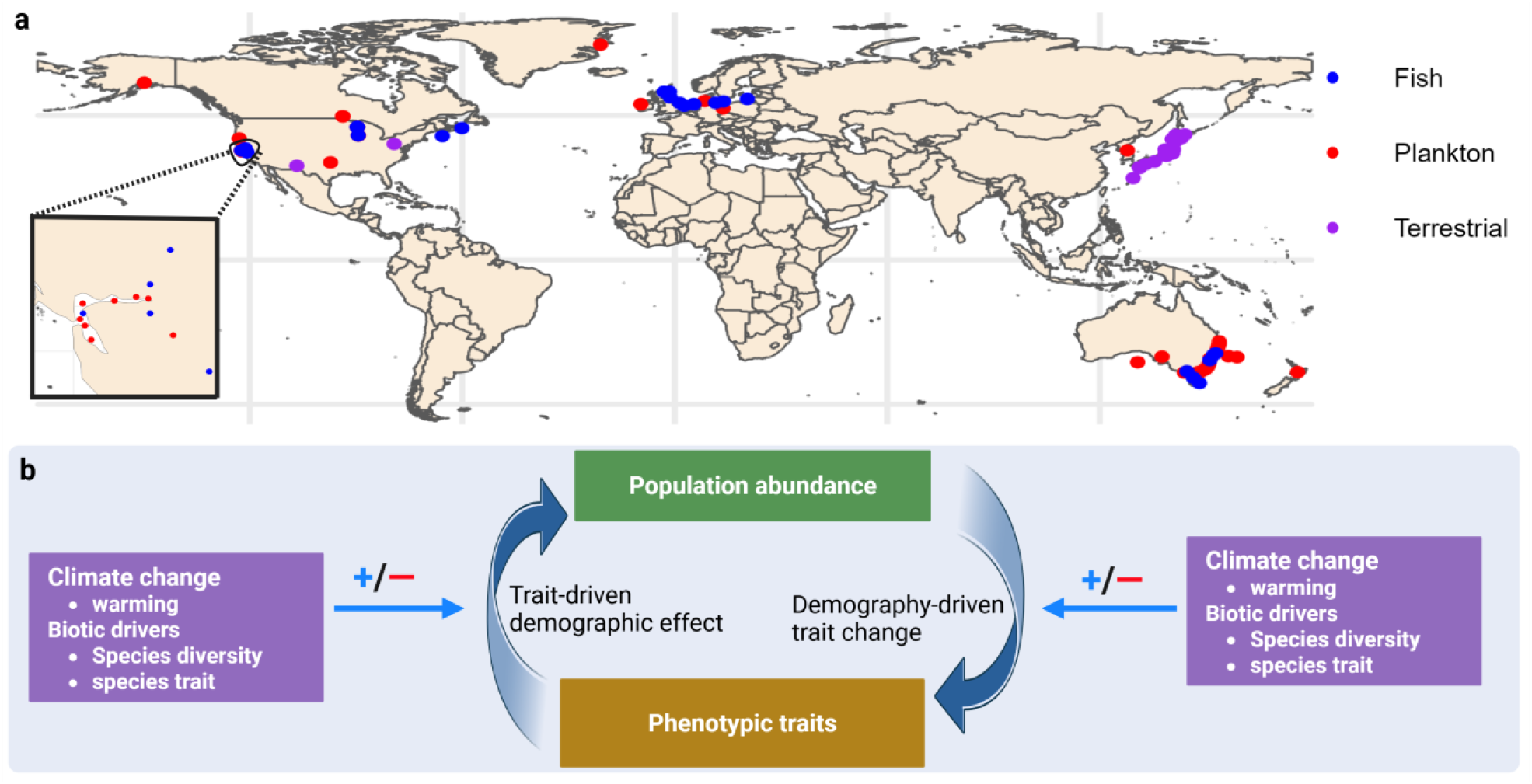
Overview of methods and sample locations. **a**, Sample locations for plankton, fish, and terrestrial species. **b**, Conceptual illustration of the eco-phenotypic feedback loop. Evolutionary and plastic shifts in phenotypic traits such as body size influence demographic processes—survival, reproduction, and competitive ability—that can ultimately alter population abundance. In turn, changes in abundance modify phenotypic traits through exerting selection pressures and inducing plastic responses. This reciprocal interaction between phenotypic traits and ecological dynamics constitutes the eco-phenotypic feedback. In this study, we investigate eco-phenotypic feedback at the level of individual focal species (e.g., *Gambusia affinis*) rather than across species.

## Methods

We compiled a long-term dataset encompassing 101 ecosystems: 51 aquatic planktonic systems monitored for 10–40 years, 30 fish systems spanning 31–47 years, and 20 terrestrial systems (beetles) spanning 9–11 years. The time series span multiple generations for each species, providing a sufficient temporal window for the phenotypic trait (body size) to change through plastic or evolutionary processes. This duration also enables the detection of concurrent shifts in abundance and body size, allowing robust assessment of eco-phenotypic feedback. We use body size as a proxy for phenotypic traits, following previous studies on eco-phenotypic feedbacks (Govaert & Klauschies 2024; Han *et al*. 2023), as body size is one of the most influential traits, exerting pervasive effects across levels of biological organization and underpinning numerous physiological, ecological, and evolutionary processes that often scale with it (Gardner *et al*. 2011; White *et al*. 2007).

### Aquatic plankton Data

Body size and abundance data were compiled from 51 long-term monitoring sites across aquatic planktonic ecosystems, comprising 20 lakes, 8 rivers, and 23 marine systems (Fig 2a). These datasets were accessed by North Temperate Lakes Long-Term Ecological Research (Magnuson *et al*. 2023a, 2024a, 2022, 2023b), Pangaea (Felden *et al*. 2023), USGS (Nejad *et al*. 2017), Greenland Ecosystem Monitoring (Christensen *et al*. 2017) and Data Observation Network for Earth (Michener *et al*. 2011). The 51 ecosystems have been consistently sampled for 10–40 years. The resident species include phytoplankton and zooplankton, with maximum generation times typically less than a week and two months, respectively (Elbrächter 1976; Gillooly 2000a; Litchman 2000). Therefore, the 10–40 years of sampling encompass numerous generations of these planktonic species. For each of the 51 aquatic planktonic ecosystems, there were three types of variables: (1) species abundance; (2) species size (i.e. body length or biovolume); (3) water temperature. For species size (2), a single metric is reported for each species and subsequently used for further analysis.

**Figure 2.**
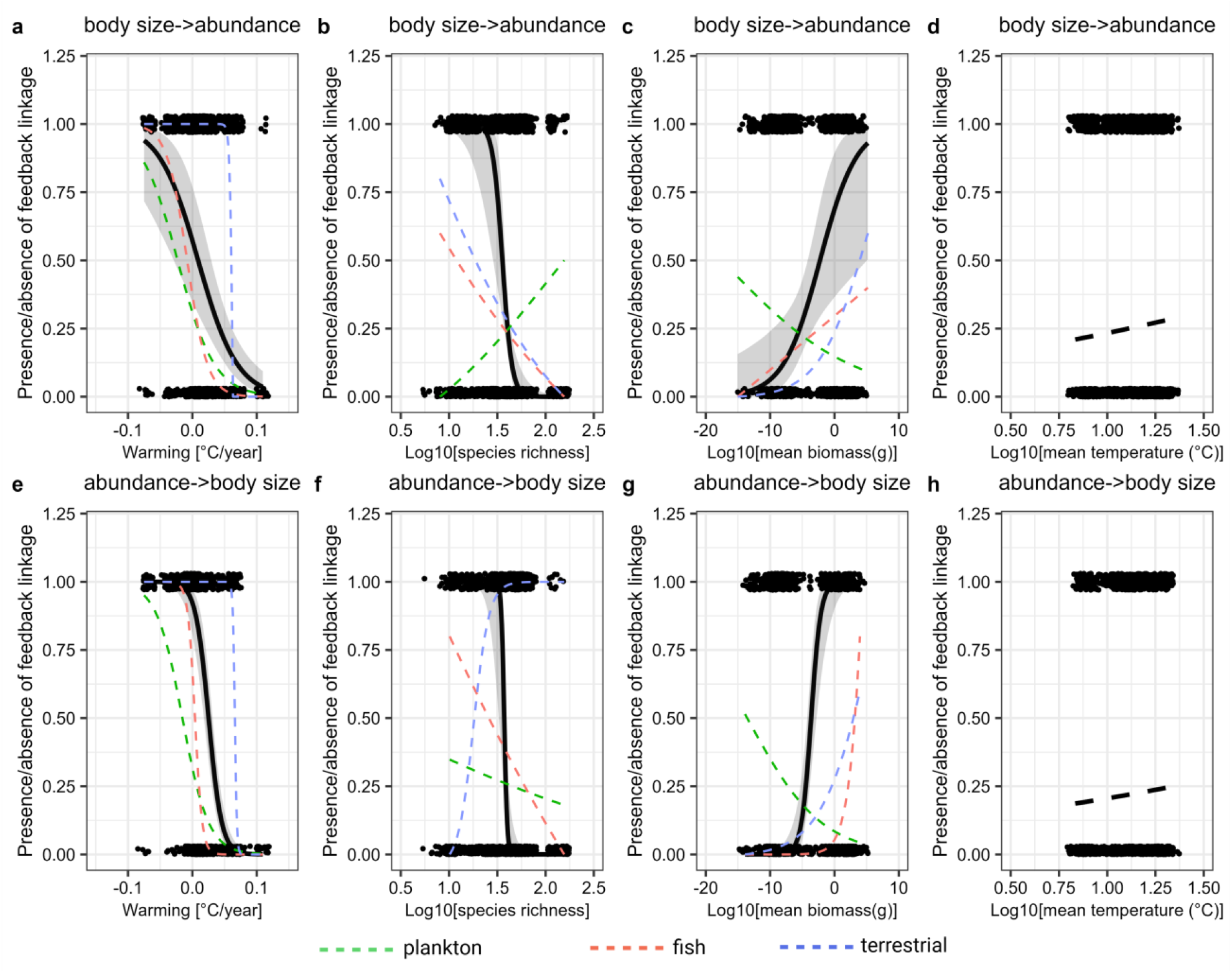
Presence/absence of eco-phenotypic feedback linkages under biotic and abiotic drivers. **a–h**, Presence (1) or absence (0) of feedback linkages in response to warming, species richness, mean species body mass and mean temperature. Link presence was identified using convergent cross mapping (CCM) and coded as 1, while absence was coded as 0. Solid black lines with shaded error bands indicate significant best-fit trendlines and corresponding 95% confidence intervals from two-sided logistic regressions across 101 ecosystems; dashed black lines indicate nonsignificant relationships. Non-bold colored dashed lines show logistic model trendlines within each taxon (plankton, fish, or terrestrial species). Points represent binary link values (0 or 1) between abundance and the phenotypic trait (body size) for each species across 101 ecosystems, with vertical jitter added to reduce overlap. Statistical results are provided in Table S1.

Next, for consistency, each species’ time series were standardized to seasonal sampling frequencies (if multiple samplings were conducted per season, e.g., monthly and bimonthly sampling data sets, those were averaged). Here, the seasonal resolution (trimonthly) is the shared consensus that can be applied to all 51 data sets, and the seasonal average here is also the most representative measure across all data sets, because the equal sampling interval is necessary for Convergent Cross Mapping (CCM) analysis. After the seasonal average, there were 0.05% missing points across the whole planktonic dataset (41 out of 85161 data points). Those missing data points were linearly interpolated using *na*.*approx* function in the package of zoo (Karakoç *et al*. 2020; Zhao *et al*. 2023). Then, the species abundance across all data sets was scaled to the same unit (individuals per litre). Species length and biovlume were standardized to the same unit (μm) and (μm^3^) respectively.

Prior to CCM analysis, we removed long‐term linear trends and seasonal cycles from each time series to avoid spurious causality from shared external drivers (Chang *et al*. 2020, 2022). Specifically, we first removed the long-term linear trend from each time series by using the residuals from a linear regression against time (Chang *et al*. 2020, 2022). Next, we removed seasonality by scaling against the mean and standard deviation of values occurring in the same year, *D*_*season*_(*t*) = (*O*(*t*) − *μ*_*season*_)/σ_*season*_, where *D*_*season*_(*t*) is the deseasoned time series, *O*(*t*) is the original time series, *μ*_*season*_ is the year mean, and the σ_*season*_ is the year standard deviation (Chang *et al*. 2020, 2022). Finally, each time series (both species abundance and size) was scaled to zero mean and unit variance.

### Fish Data

Fish size and abundance data were compiled from 30 monitoring sites, including six rivers, five lakes and 19 marines. These datasets were accessed by North Temperate Lakes Long-Term Ecological Research (Magnuson *et al*. 2024c, b), International Council for the Exploration of the Sea (ICES) (Fish trawl surveys 2025), and Reef Life Survey (Edgar & Stuart-Smith 2014). These ecosystems have been consistently sampled for 31–47 years. Again, for each Fish ecosystem, there were three types of variables: (1) species abundance; (2) species size (i.e. body length); (3) water temperature.

For consistency each species time series of abundance and size were standardized to the yearly sampling frequency (if multiple samplings were conducted per year, e.g., quarterly sampling data sets, those were averaged). Again, the yearly resolution is shared across all data sets. After the yearly average, there were no missing points across the fish dataset. Then, the fish abundance across all data sets was standardized to the same unit (caught per effort). Species length was standardized to the same unit (mm). As all fish data were standardized to yearly intervals, there was no need to remove long-term linear or seasonal trends from each time series. Therefore, the original data were utilized for further analysis. Finally, each time series (both species abundance and size) were scaled to zero mean and unit variance.

### Terrestrial species

Terrestrial species size and abundance data were compiled from 20 monitoring sites, all focused on ground beetles. All data were retrieved from BioTIME (Dornelas *et al*. 2018). These ground beetles were consistently sampled for 10–11 years. Given a maximum generation time of typically smaller than a year, the sampling period encompasses numerous generations across all species (Kirk 1975; Matalin 2007; Rivard 1964). Again, for each site, there were three types of variables: (1) species abundance; (2) species size (i.e. mg); (3) surface temperature. Each site reports both abundance and size. The daily temperature and precipitation for each site were retrieved from NOAA’s National Centers for Environmental Information (GHCN-Daily, Version 3 2012).

Next, to ensure consistency, each species’ time series were standardized to seasonal sampling frequencies (if multiple samplings were conducted per season, those were averaged). After calculating seasonal averages, 0.52% of data points were missing in the dataset. These missing values were subsequently linearly interpolated using the *na*.*approx* function from the zoo package (Karakoç *et al*. 2020; Zhao *et al*. 2023). Then, long-term linear and seasonal trends were removed from each time series using the same methodology applied to the plankton data. Finally, each time series, including both species abundance and size, was standardized to have a mean of zero and a unit variance.

### Presence/absence of eco-phenotypic feedback linkages under biotic and abiotic drivers

For each species (plankton, fish or terrestrial species), we identified the presence or absence of the feedback linkages (i.e. size →abundance, abundance →size) using CCM (Fig 1). CCM is grounded in Takens’s theorem for dynamic systems, which infers causal relationships among variables from empirical time series (Deyle & Sugihara 2011; Sugihara *et al*. 2012). If a causal variable X (e.g., species abundance) and a response variable Y (e.g., body size) belong to the same dynamical system, information about X is embedded in Y. Thus, the state of X can be predicted from Y through a process known as cross mapping (cross-prediction) (Sugihara *et al*. 2012). CCM quantifies causality by assessing the degree to which X leaves a signature in the time series of Y. In this study, the appropriate embedding dimensions for cross-mapping E were determined by the optimized the hindcast crossmapping in which X(t) projected one-step backward to Y(t-1), examining values of E from 2 to square root of *n*, where n is the length of the time series (Karakoç *et al*. 2020). Thus, we computed *n* as the geometric mean time series length across all data sets. E was finally examined from 2 to 7 across all data sets. We used simplex projection to select the best E that gave the highest prediction skill (Hsieh *et al*. 2005).

To accommodate the fact that the causal variables (e.g., species abundance) can exhibit time-delayed effects on the affected variables (e.g., species size), we carried out 0 to 1 year (i.e. 0~4 time point for plankton and terrestrial species but 1 time point for fish) lagged CCM analyses, in which we retained the CCM with the time lag resulting in the highest cross mapping skill ρ (Ye *et al*. 2019). The criterion for testing causality is the convergence toward higher cross-mapping skill with increasing library length (i.e., the number of points used for state space reconstruction) (Sugihara *et al*. 2012). An increase in library length enhances the density of points in the reconstructed attractor, facilitating more precise identification of nearest neighboring points for predictions. This, in turn, leads to improved predictive accuracy (Sugihara & May 1990). Convergence can be assessed by evaluating whether there is a significant monotonic increasing trend in cross-mapping skill ρ as library length increases using Kendall’s τ test (Chang *et al*. 2020), and whether the ρ at the largest library length is significantly greater than that the ρ at the smallest library length using Fisher’s Z test (Chang *et al*. 2020). In this study, library length was set from the minimum (E) to the maximum length (i.e., the length of the entire time series). Throughout this study, an interaction link (e.g., X→Y) was considered causal if both Kendall’s τ test and Fisher’s Z test were statistically significant (*P* < 0.05) for testing convergence (Chang *et al*. 2020).

Then, the warming rate, representing the long-term warming trend of water temperature or land surface temperature in each ecosystem, was measured using the Theil–Sen median-based trend estimator, implemented via ‘sens.slope’ from ‘trend’ package (Chang *et al*. 2020; Mohsin & Gough 2010). This method was chosen due to its widespread use in quantifying warming rates (Pinsky *et al*. 2025; Tong *et al*. 2023), its robustness to episodic extreme events, and its ability to provide stable estimates of long-term trends in the presence of outliers (Mohsin & Gough 2010). Species mean body mass was computed as average across whole time series, estimated from biovolume, assuming that organisms have the same density as water, if unavailable, was obtained from published sources (Cohen & Lough 1981; Huntley 1992). Similarly, mean temperature was computed as average across whole time series. Species richness for each of the 101 ecosystems was calculated as the total number of species present within each system.

Next, we employed a generalized linear mixed model (glmmTMB function from the glmmTMB package (Brooks *et al*. 2017)) with a binomial error distribution to test the main effects of warming, mean temperature, species richness, mean species body mass on presence/absence of the feedback linkages—either from phenotypic trait (body size) to abundance (size →abundance) or from abundance to body size (abundance → size) (table S1). To ensure normality, mean temperature, species richness, and mean species body mass were log-transformed using base 10. Each of these predictors in the model were normalized using the scale function from base R. The response variable in this model was binary, with 1 indicating the presence of a feedback linkage identified using CCM and 0 indicating its absence. Last, we set species groups (e.g. plankton) nested within sample locations (e.g. Trout Lake) as random intercepts. Details of statistics see Table S1.

### Strength of eco-phenotypic feedback linkages under biotic and abiotic drivers

The strength of feedback linkages (i.e. size →abundance or abundance →size) was quantified by CCM based on cross-mapping skill ρ (i.e., correlation coefficient between observations and CCM predictions) using the maximal training set (Sugihara *et al*. 2012). To ensure comparability across 101 ecosystems, we accounted for variation in cross-map skill ρ arising from differences level of noise in time-series observed in each ecosystem. Following (Chang *et al*. 2020), linkage strength was therefore standardized by dividing each value by the maximum cross-map ρ within a system, yielding scaled strengths between 0 and 1 that reflect the relative importance of each causal link (Chang *et al*. 2022).

Next, we employed a generalized linear mixed model using the glmmTMB function from the *glmmTMB* package (Brooks *et al*. 2017) with a Gaussian error distribution to test the main effects of warming, mean temperature, species richness, and mean species body mass on linkage strength—either from phenotypic traits (body size) to abundance (size → abundance) or from abundance to body size (abundance → size) (Table S2). Again, to ensure normality, mean temperature, species richness, and mean species body mass were log10-transformed, and all of these predictors were normalized using the scale function in base R. Species groups (e.g., plankton) nested within sampling locations (e.g., Trout Lake) were included as random intercepts. Full statistical results are provided in Table S2.

All analyses were done with R (ver. 4.1.2). The CCM analysis was examined using the rEDM package (version 1.2.3).

## Results

We first used a binary response variable to indicate the presence or absence of feedback linkages between the phenotypic trait (body size) and population abundance, with links identified by Convergent Cross Mapping (CCM). Across 101 ecological systems monitored for 10 to 47 years, our analysis showed that warming weakened eco-phenotypic feedback loop, as evidenced by a negative association between warming and the proportion of both linkages (abundance → size and size → abundance) (Fig 2a, 2e). Species richness likewise weakened the feedback loop (Fig 2b, 2f), whereas mean species body mass strengthened it (Fig 2c, 2g). However, mean temperature showed no effect on either linkage of abundance → size or size → abundance (Fig 2d, 2h).

Within taxonomic groups (plankton, fish, or terrestrial species), the effects of species richness and mean species body mass on eco-phenotypic feedback loops were taxon-specific, whereas warming consistently weakened feedback strength across all taxa (Fig 2). In most taxa, species richness weakened the eco-phenotypic feedback loop (Fig 2b, 2f), whereas mean species body mass strengthened it (Fig 2c, 2g).

Last, we used the strength of feedback linkages, as quantified by CCM, as the response variable. Across 101 ecological systems, mean temperature only increased the strength of the size → abundance linkage (Fig 3d). In contrast, warming, species richness, and mean species body mass had no effects on the strength of either linkage (abundance → size or size → abundance) (Fig 3). Taken together, these results indicate that warming and species richness weakened eco-phenotypic feedback loops by reducing the proportion of both linkage types rather than altering their strength, whereas mean species body mass strengthened the feedback loop by increasing them.

**Figure 3.**
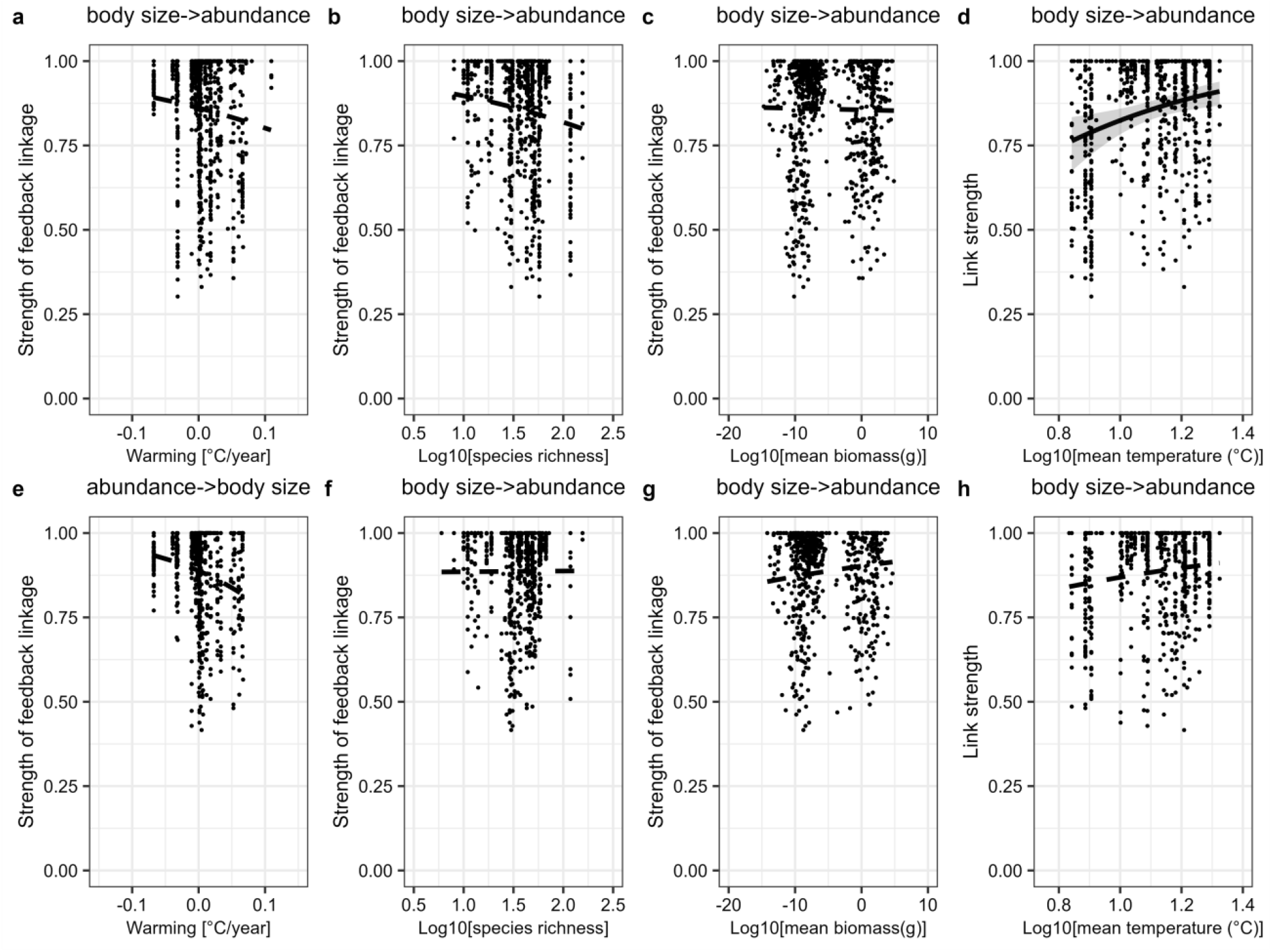
Strength of eco-phenotypic feedback linkages under biotic and abiotic drivers. **a–h**, Strength of feedback linkages in response to warming, species richness, mean species body mass and mean temperature. Link strength was quantified using convergent cross mapping (CCM), with larger values indicating stronger feedback linkages. Solid black lines with shaded error bands indicate significant best-fit trendlines and corresponding 95% confidence intervals from two-sided regressions across 101 ecosystems; dashed black lines indicate nonsignificant relationships. Statistical results are provided in Table S2.

## Discussion

Our study provides the first systematic evaluation of eco-phenotypic feedback loop in response to climate warming and biotic drivers. Using long-term field data from 101 ecosystems—each spanning multiple generations per species and providing sufficient temporal durations to detect changes in both phenotypic traits and population abundance—we found that warming and species richness generally weakened eco-phenotypic feedback loops, whereas mean species body size strengthened them. These effects were driven by changes in the proportion of both linkage types (abundance → size or size → abundance).

The phenotypic trait of body size respond rapidly to environmental change through phenotypic plasticity or evolutionary adaptation (Merckx *et al*. 2018) and exerts a substantial influence on population abundance (Martins *et al*. 2023). Previous experimental studies have documented that the ecophenotypic feedback are tightly linked in single-species populations (Gibert *et al*. 2022, 2023), a two-species competitive interactions (Govaert & Klauschies 2024), and a microbial food web (Han *et al*. 2023). Our study further showed that warming tended to weaken the feedback loop (Fig 2).

Previous evidence has documented weaker eco-phenotypic feedback at higher temperatures from an experimental protist food web (Han *et al*. 2023). This may be attributed to the differing response rates of phenotypic trait of body size and abundance to warming (Genner *et al*. 2010). Higher temperatures can lead to a significant reduction in species body size (Audzijonyte *et al*. 2020; Terry *et al*. 2022), while it can have slight effect on population abundance (Sebastian *et al*. 2012). Alternatively, higher temperatures have shown to accelerate growth, leading to an rapid increase in population abundance (Brandenburg *et al*. 2019; Han *et al*. 2023), while simultaneously either leaving body size unaffected or slightly altered body size (Crozier *et al*. 2010; Genner *et al*. 2010). These different responses rate among body size and abundance may weaken the feedback between them (Han *et al*. 2023).

We found that higher species richness also weakens eco-phenotypic feedback loop. Previous experimental studies have similarly shown that increased species richness reduces the proportion of links from phenotypic traits body size to abundance (body size→ abundance) (Govaert & Klauschies 2024). Here, enhanced species richness can subject organisms to a broader spectrum of ecological interactions, including interspecific competition and predation, which may disrupt the feedback between size and abundance (Govaert & Klauschies 2024). Evidence has documented that adding interspecific competition and predation can weaken the link strength between them (Govaert & Klauschies 2024; Han *et al*. 2023).

We found that mean species body size strengthens the eco-phenotypic feedback loop. To date, no direct empirical evidence has been documented to validate this trend. Species with larger body mass exhibit longer generation times and slower population turnover (Gillooly 2000b; White *et al*. 2007), allowing both body size and population abundance to respond more gradually to environmental changes. This extended adaptation period may contribute to a more stable and synchronized relationship between body size and population abundance over time.

In summary, by using long-term field data covering multiple generations for each species, our findings show that climate warming weakens ecophenotypic feedback loops, highlighting their vulnerability to anthropogenic pressures. Understanding how eco-phenotypic feedback responds to environmental change may help predict trait and population abundance dynamics in a warming world.

## Author Contributions

Q.H.Z., F.D.L. and Y.X.G.W designed the research. Q.H.Z. and Y.X.G.W performed the study and analyzed the data. Q.H.Z and G.S performed the CCM analysis. Q.H.Z., F.D.L. and Y.X.G.W led the writing, with contributions fromC.X. and G.S.

## Data and code availability

The data reproducing all the results in this study are available on GitHub (https://github.com/QZhao16/link.abundance-size).

## Acknowledgements

We thank all participants in each long-term monitoring site. Q.Z. acknowledges funding from the concerted research action (ARC) from the special research fund from the University of Namur, Belgium (ARC grant DIVERCE, Convention 18/23-095).

## Supplementary Tables

**Table S1.** Presence/absence of eco-phenotypic feedback linkages under biotic and abiotic drivers. A generalized linear mixed model (glmmTMB function in the *glmmTMB* package) with a binomial error distribution was used to test the main effects of warming, mean temperature, species richness, and mean species body mass on the presence/absence of feedback linkages (size → abundance or abundance → size). The response variable was binary, with 1 indicating the presence of a feedback linkage identified using convergent cross mapping (CCM) and 0 indicating its absence. Species groups (e.g., fish) nested within sampling locations (e.g., Trout Lake) were included as random intercepts. Mean temperature, species richness, and mean species body mass were log10-transformed to improve model performance. \*\*\**P*< 0.001, \*\**P*<0.01, \**P*<0.05.

|  | presence/absence of body<br>size → abundance |  |  | presence/absence of<br>abundance → body size |  |  |
| --- | --- | --- | --- | --- | --- | --- |
|  | coefficient | Z | P value | coefficient | Z | P value |
| (Intercept) | -1.10 | -4.95 | **<8e-8\*\*\*** | -1.27 | -11.92 | **<2e-16\*\*\*** |
| Species richness | -0.99 | -3.31 | **<0.0001\*\*\*** | -3.44 | -9.45 | **<2e-16\*\*\*** |
| Warming | -0.99 | -4.65 | **<4e-6\*\*\*** | -2.87 | -11.72 | **<2e-16\*\*\*** |
| Mean body mass | 0.81 | 2.66 | **0.008\*\*** | 3.10 | 8.59 | **<2e-16\*\*\*** |
| Mean Temperature | 0.12 | 1.09 | 0.28 | 0.11 | 1.12 | 0.26 |

**Table S2.** Strength of eco-phenotypic feedback linkages under biotic and abiotic drivers. A generalized linear mixed model (glmmTMB function in the *glmmTMB* package) with a gaussian error distribution was used to test the main effects of warming, mean temperature, species richness, and mean species body mass on the strength of feedback linkages (size → abundance or abundance → size). Strength of feedback linkages was quantified using convergent cross mapping (CCM), with larger values indicating stronger feedback linkages. Species groups (e.g., fish) nested within sampling locations (e.g., Trout Lake) were included as random intercepts. Mean temperature, species richness, and mean species body mass were log10-transformed to improve model performance. \*\*\**P*< 0.001, \*\**P*<0.01, \**P*<0.05.

|  | Strength of body size → abundance |  |  | Strength of abundance → body size |  |  |
| --- | --- | --- | --- | --- | --- | --- |
|  | coefficient | Z | P value | coefficient | Z | P value |
| (Intercept) | 1.80 | 15.75 | **<2e-16\*\*\*** | 2.06 | 16.60 | **<2e-16** |
| Species richness | -0.17 | -1.53 | 0.13 | 0.01 | 0.05 | 0.96 |
| Warming | -0.14 | -1.04 | 0.30 | -0.25 | -1.61 | 0.11 |
| Mean body mass | -0.02 | -0.14 | 0.89 | 0.15 | 1.04 | 0.30 |
| Mean Temperature | 0.33 | 3.00 | **0.002\*\*** | 0.17 | 1.40 | 0.16 |

